# Pharmacophore-based peptide biologics neutralize SARS-CoV-2 S1 and deter S1-ACE2 interaction *in vitro*

**DOI:** 10.1101/2020.12.30.424801

**Authors:** Masaud Shah, Sung Ung Moon, Hyun Goo Woo

**Author notes:** Contributed equally (Masaud Shah, Sung Ung Moon). **Corresponding author**: Hyun Goo Woo, M.D., Ph.D.

## Abstract

Effective therapeutics and stable vaccine are the urgent need of the day to combat COVID-19 pandemic. SARS-CoV-2 spike protein has a pivotal role in cell-entry and host immune response, thus regarded as potential drug- and vaccine-target. As the virus utilizes the S1 domain of spike to initiate cell-attachment and S2 domain for membrane fusion, several attempts have been made to design viral-receptor and viral-fusion blockers. Here, by deploying interactive structure-based design and pharmacophore-based approaches, we designed short and stable peptide-biologics *i.e*. CoV-spike-neutralizing peptides (CSNPs) including CSNP1, CSNP2, CSNP3, CSNP4. We could demonstrate in cell culture experiments that CSNP2 binds to S1 at submicromolar concentration and abrogates the S1-hACE2 interaction. CSNP3, a modified and downsized form of CSNP2, could neither interfere with the S1-hACE2 interaction nor bind to S1. CSNP4 exhibited dose-dependent binding to both S1 and hACE2 and abolished the S1-hACE2 interaction *in vitro*. CSNP4 possibly enhance the mAb-based S1 neutralization by limiting the spontaneous movement of spike receptor-binding domain (RBD), whereas CSNP2 allowed RBD-mAb binding without any steric hindrance. Taken together, we suggest that CSNP2 and CSNP4 are potent and stable candidate peptides that can neutralize the SARS-CoV-2 spike and possibly pose the virus to host immune surveillance.

## Background

Since its appearance in Dec 2019, COVID-19 has crippled the world economy and affected almost every aspect of life. According to the WHO, ~82.5 million people have been infected and ~1.8 million lives have been claimed by COVID-19, by the end of year 2020. The shared research data contribution and innovative vaccine development ideas by the scientists around the world, led to the development of dozens of mRNA vaccine, two of which, mRNA-1273 and BNT162b2, are approved by FDA (1, 2) on emergency basis and many are undergoing clinical trials (3). Sputnik V vaccine developed by Gamaleya Research Institute” has been approved by the Ministry of Health of the Russian Federationa and AZD1222, developed by AstraZeneca at Oxford University, has been recently authorized by the UK Medicines and Healthcare Products Regulatory Agency (MHRA). The two vaccine candidates developed by Pfizer and BioNTech (BNT162b2) and Moderna (mRNA-1273) have shown ~95% efficacy in their phase III clinical trials (1, 2). The immune evasion-strategies utilized by SARS-CoV-2 make the vaccine and drug development processes more difficult (4–6). SARS-CoV-2 exploits both structural and non-structural proteins to perturb host immune response. Eight out of the 23 proteins of SARS-CoV-2 suppressed the induction of type-1 interferon by Sendai virus, whereas S and NSP2 exhibited IFN induction (7). SARS-CoV-2 Nsp1 binds to the host ribosome (40S) and obstruct the mRNA entry, thereby blocking the RIG-I-dependent IFN response (8). Many factors including mutation of the virus, halflife of the neutralizing antibodies, safety of vaccine, and above all, the availability of the vaccine is of priority concerns in this scenario. The capacity of host adaptation by SARS-CoV-2 as exemplified by the previous D614G mutation (9) and recent N501Y, D796H mutations in spike are alarming and raised concerns over the efficacy of neutralizing antibodies response induced by these vaccines that utilize wild type spike antigen (10).

Receptor recognition and cell-entry of the SARS-CoV-2 are requisite and hold high priority for designing effective therapeutic interventions. The spike glycoprotein (S) is expressed as a trimeric complex on the surface of virus, containing two subdomain S1 and S2 (11). The S1 domains contains receptor, angiotensin-converting enzyme 2 (ACE2), binding domain (RBD), whereas the S2 domain facilitates membrane fusion after the enzymatic cleavage of the S1 domain. The host-dependent protease activation of the SARS-CoV-2 entry is a crucial determinant of its infectivity and pathogenesis (12). Lysosomal proteases, including cathepsins and cell-surface expressed type II transmembrane serine proteases (TMPRSS2) play important roles in viral activation, exhibiting an augmented effect on SARS-CoV-2 entry in the presence of furin (13). This structural changes in the S protein allows the S2 domain for host-cell membrane fusion while the ACE2-bound S1 domain sheds in the extracellular environment (14).

In the trimeric spike, the RBD switches between “up” and “down” conformations to facilitate ACE2 binding and evade immune surveillance, respectively (11, 15). Masking of the RBD can lead to the paradox of immune evasion with less infectivity of the virus. The D614G mutation in the spike protein, is considered to have increased the “up” conformation and the overall density of the spike protein at the surface of the virus. This makes the SARS-CoV-2^D614G^ more infectious and more sensitive to the neutralizing antibodies (14, 16). Hence, the spontaneous conformation-switching of the RBD represents a major challenge to develop neutralizing antibodies and vaccines.

Detailed structural and protein-protein interfacial (PPI) insights in ACE2-RBD are fundamental to design effective therapeutic interventions against SARS-CoV-2 (17, 18). PPI information and decoy approaches have been used to design structurally constrained therapeutic peptides that are able to retain their structures and efficiently grab the shallow surfaces at targets (19–21). A peptide, S471-503, derived from the ACE2 binding region of the SARS-CoV RBD hinders the ACE2-RBD interaction and viral entry into the cell (22). Another peptide against SARS-CoV constructed by the glycine linkage of two separate segments (a.a. 22-44 and 351-357) of ACE2 also exhibited efficient antiviral activity (IC50 = 0.1 μM) (23). Corroborating this approach, a truncated 23-mer (a.a. 21-43) peptide (SBP1) from the α1 helix of the ACE2 exhibited SARS-CoV-2 glycosylated-RBD-binding in nano-molar concentration (K_D_=47 nM) (24). However, this binding was only observed in the insect-derived RBD, and the peptides did not show any binding to HEK cell-expressed or other insect-derived SARS-CoV-2-RBD variants. This imply that the 351-357 region contains pharmacophores essential for the ACE2-RBD interaction. To further support the idea of peptide antidotes against SARS-CoVs, Shuai et al. has reported a peptide EK1, derived from the HR2 motif of the S2 domain of the SARS-CoV (25). Modification of this peptide with some lipid and polyethylene glycol (PEG) could inhibit SARS-CoV-2-spike as well as host-cell fusion, therefore, considered as pan-coronavirus fusion inhibitor (26).

Recently, by collecting PPI knowledge and understanding the dynamics of the complex protein interactions, we have designed effective therapeutic peptides (27–29). In the current study, we attempted similar approaches to design structurally constrained peptides, which could address the above-mentioned paradoxes of the spontaneous RBD-conformation switching and the interaction of S1 with soluble and membrane bound ACE2. We and others have delineated the ACE2-RBD interface and identified key residues that contribute to the binding strength of ACE2-RBD (15, 17, 18, 30). Point mutation analyses could confirm that some of these residues are vital to the ACE2-RBD interface (31). Collecting together these data, we could identify pharmacophores on both ACE2 and RBD that are crucial to their binding. Taking advantage of the possibility that some residues in the α1 helix of the ACE2 are sub-optimal for the RBD-binding (32), these residues can be substituted to augment the ACE2-RBD binding. In consideration of the crucial pharmacophores/hotspots and suboptimal residues, we designed short CoV-spike-neutralizing peptides (CSNPs) and demonstrated their effects on the S1-hACE2 binding *in vitro*.

## Results

### Design of the candidate CSNPs

We set up two strategies to design CSNPs to neutralize the SARS-CoV-2 spike as well as address the dilemma of reversible position-switching of the RBD and the masking of S1 by soluble ACE2. First, the main scaffold of the α1 helix of ACE2, which is mainly involved in RBD-interaction, was extracted as a starting structure to design and assemble helical CSNPs (CSNP1-3). Second, the ACE2-interacting motifs of RBD were extracted and assembled into CSNP4 to restrict the RBD movement and block its binding to ACE2.

ACE2 mainly utilizes polar and charged residues in its α1 helix to grab the RBD in its “up” conformation (15) through salt-bridges and hydrogen bonds (**Table 1**). The first three amino acids, Ile21, Glu22, and Glu23 in α1, are exposed to solvent without involving in the RBD binding; nonetheless, these residues are attributable to establish the α1 helix. Five residues, Gln24, Asp30, Lys31, Asp38, and Tyr41 were found as the major hotspots from α1 that contribute the highest binding energy to the ACE2-RBD complex (**Supplementary Figure 1A, Table 1**). In addition to α1 helix, ACE2 utilize Lys353 to anchor to the RBD of both SARS-CoV and SARS-CoV-2, and shares the second highest binding energy among the ACE2-RBD interface residues **(Table 1)**. As the Lys353 lays at the hinge of β3-β4 that are stapled by a disulfide bond between Cys344 and Cys361, the flexibility and freedom of this amino acid might be restricted. This allows the hydrogen-bond-network intact between ACE2 (Lys353) and RBD (Gly496, Gln498, and Gly502) (**Figure 1A, Table 1**). Together, we identified five pharmacophores i.e. Asp30, Lys31, Asp38, and Tyr41 in α1 and Lys353 in β3-β4 that keep the ACE2-RBD intact (for simplicity four of them are shown in surface electrostatic map in figure 1A). The COOH^-^ group of Asp30 serves as hydrogen bond acceptor, which is in the vicinity of NH3^+^ group of Lys417 of RBD. The NH3^+^ group of Lys31 of α1 lays between the Glu35 of α1 and Glu484 of RBD and establishes salt bridge, alternatively. The COOH^-^ group of Asp38 in α1 is indispensable for the stability of Lys353 in ACE2 and it also makes crucial contacts with Tyr449 and Gln498 of the RBD. The bulky side chains of Tyr41 occupies the hydrophobic space between the a.a. 350-359 segment and the N-terminus of a.a. 21-46 segment in peptide. Besides, the Tyr41-Thr500 hydrogen bond between ACE2 and RBD restricts the rotation of Tyr41. The NH3^+^ group of Lys353 is very crucial pharmacophore with respect to the ACE2-RBD interaction (**Table 1**).

**Figure 1:**
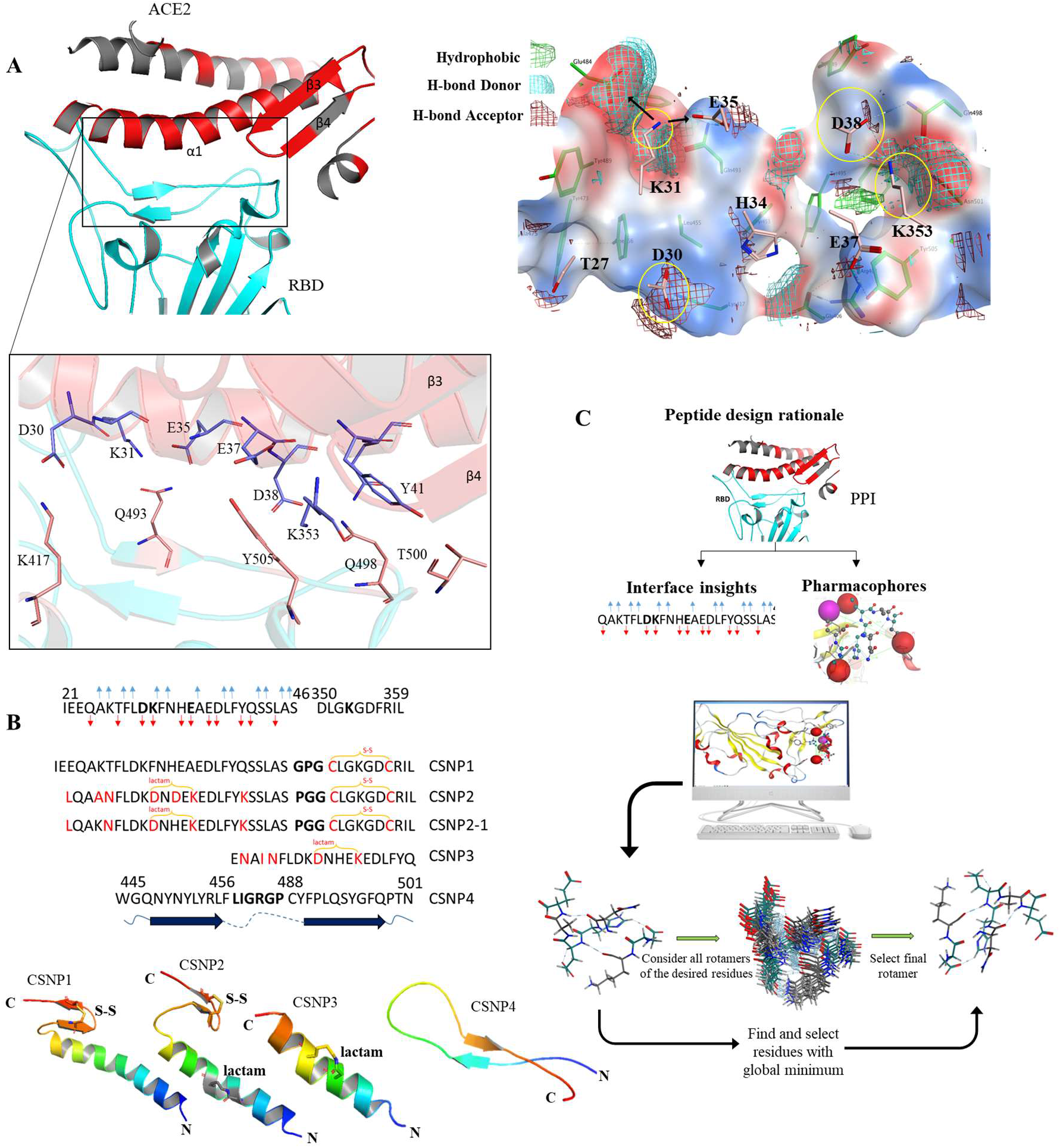
The SARS-CoV-2 Spike RBD-hACE2 interface analyses and CSNPs peptides designing rationale. **A**) Identification of the crucial residues at the ACE2-RBD interface using interactive structures and surfaces strategy. The smooth surfaces (blue, +ve; red, −ve; white, neutral) are drawn over the RBD residues while the meshes represents the electrostatic surface around the interface residues of ACE2. **B**) The sequence and secondary structure representation of the designed CSNPs. **C**) The overall protocol of CSNPs design. The interface residues were identified, vetted for their energy contribution at the interface (Supplementary figure 1), and featured for pharmacophores or non-pharmacophores. The potential residues were substituted to augment the ACE2-RBD binding.

**Table 1:** The Hydrogen bonds at the interface of ACE2-RBD and their bonds-strength in term of energies.

| ACE2 | RBD | E<sup>kcal/mol</sup> | Dist Å |
| --- | --- | --- | --- |
| Asp30 | Lys417 | -33.5 | 2.71 |
| Lys353 | G1y496 | -11.1 | 2.71 |
| Lys353 | G1n498 | -7.3 | 2.68 |
| Lys31 | G1n493 | -6.3 | 2.93 |
| Lys31 | G1u484 | -5.64 | 3.24 |
| G1u35 | G1n493 | -4.9 | 2.71 |
| Lys353 | G1y502 | -4.9 | 2.75 |
| G1u37 | Tyr505 | -2.8 | 2.65 |
| G1n24 | Asn487 | -1.9 | 2.81 |
| Tyr83 | Asn487 | -1.7 | 2.64 |
| G1n42 | G1y446 | -1.6 | 2.92 |
| Asp38 | G1n498 | -1.2 | 3.57 |
| Lys31 | Tyr489 | -0.5 | 4.35 |

For the helical peptides, the scaffold of α1 helix (a.a. 21-46) of ACE2 was truncated and linked with the β3-β4 (a.a. 350-359) through a Gly-Pro-Gly (GPG) linker. The freedom of Lys353 was restricted by creating an S-S bond between position D350C and F356C. We designed this peptide as a parent peptide (CSNP1, **Figure 1B**). Next, the complementarity of the electrostatic surface was examined, revealing potential points that could enhance the binding affinity between CSNPs and RBD. Mutations of the potential residues were constructed with all possible permutation under the consideration of the available volume, surface complementarity, total binding energy, and stability of the complex. The resulting peptides database (81 mutants) with single substitutions and their respective binding-affinities and binding stabilities were recorded and utilized in next round of residues scan. The top five substitutions of each residue (*i.e*., Glu23, Lys26, Thr27, His34, Gln42) were selected and implemented in multi-substitution peptides construction step. Monitoring of their binding energies could identify the CSNP2 and CSNP2-1 as the best fit peptides to the RBD interface (**Supplementary Table 1**). To retain their helicity, a structural constraint (lactam bridge) was created between the side chains of the non-interface residues Phe32Asp and Ala36Lys (**Figure 1B**). The GPG linker was changed to PGG in CNSP2 to enhance the flexibility of the loop. A shorter constrained peptide, CSNP3, was constructed by considering the pharmacophores of α1 helix to validate both the importance of Lys353 and self-sufficiency of α1 helix for RBD binding.

CSNP4 was designed under the consideration of the spontaneous position switching as well as ACE2-RBD interface residues of the RBD. Two amino acid stretches of 445-456 and 488-501 which participate in ACE2 binding, were truncated from the RBD and joined through flexible linker, LIGRGP, to optimally orient the joining peptides and retain its target-binding ability. In its resting (RBD^down^) position, the same sheet-loop-sheet motif lays between the NTD and the RBD domains of the adjustment S protomer, as we have shown previously (30). Thus, in addition to ACE2-RBD hindrance, we suggest that CSNP4 can present RBD for the immune surveillance by limiting its spontaneous position switching. The overall strategy for peptide designing is outlined in **Figure 1C**.

### Structural-constrains stabilize the CSNPs

Structural stability and resistance to enzymatic degradation are important features in designing small therapeutic peptides. Moreover, fold-on-binding requires time and often peptides lose their target specificity if the structures are not intact (28); therefore, the structures of CSNP1-3 were stabilized by applying structural restrains, and the two amino acid stretches of the CSNP4 were joined through a shorter loop “LIGRGP”. To check their structural stability, these peptides were simulated in an aqueous environment as a function of time. To validate the stability, we also simulated SBP1, a previously reported structurally unrestrained ACE2-derived RBD-binding peptide. Overall, there was a considerable root mean square deviation (RMSD) in all atoms of the peptides in the 2^nd^ and 3^rd^ quarter of the simulation (**Figure 2A**). To track the fluctuation of RMSD at atomic level, the root mean square fluctuation (RMSF) of all atoms of all five peptides were calculated. The terminal atoms of all peptides, particularly CSNP4, exhibited considerable fluctuation as compared to the atoms in peptide bodies. However, CSNP1 and SBP1 exhibited high fluctuation in atoms ranging from 100-170, as compared to CSNP2 and CSNP3 (**Figure 2B**). The radius if gyration (Rg), which predicts the folding with compactness of the peptides, showing that SBP1 and CSNP3 undergoes a dramatic shift and exhibit a shrinkage in structure (**Figure 2C**). This suggests that the hydrogen bonds between the sidechains of these peptides have probably acquired a new pattern and perhaps disordered the peptides structure. To validate these data, 1000 structural frames were extracted from the 200 ns MD trajectory of each peptide and inspected in their secondary structural alteration. The 3D motions and changes in their secondary structure, as a function of time, were preserved in 3D animation (**Figure 2D and Supplementary movie 1**). Remarkably, CSNP1 and SBP1 were shifted from helical to an irregular looped structures, permanently losing their structural helicity; however, CSNP3 partly retained its helical structure. Among the helical peptides CSNP2 held its structure intact; however, its C-terminal S-S constrain region remained flexible at the PGG junction (**Supplementary movie 1**). These data suggest that constrains stabilize peptide structures and could possibly retain their target binding affinity.

**Figure 2:**
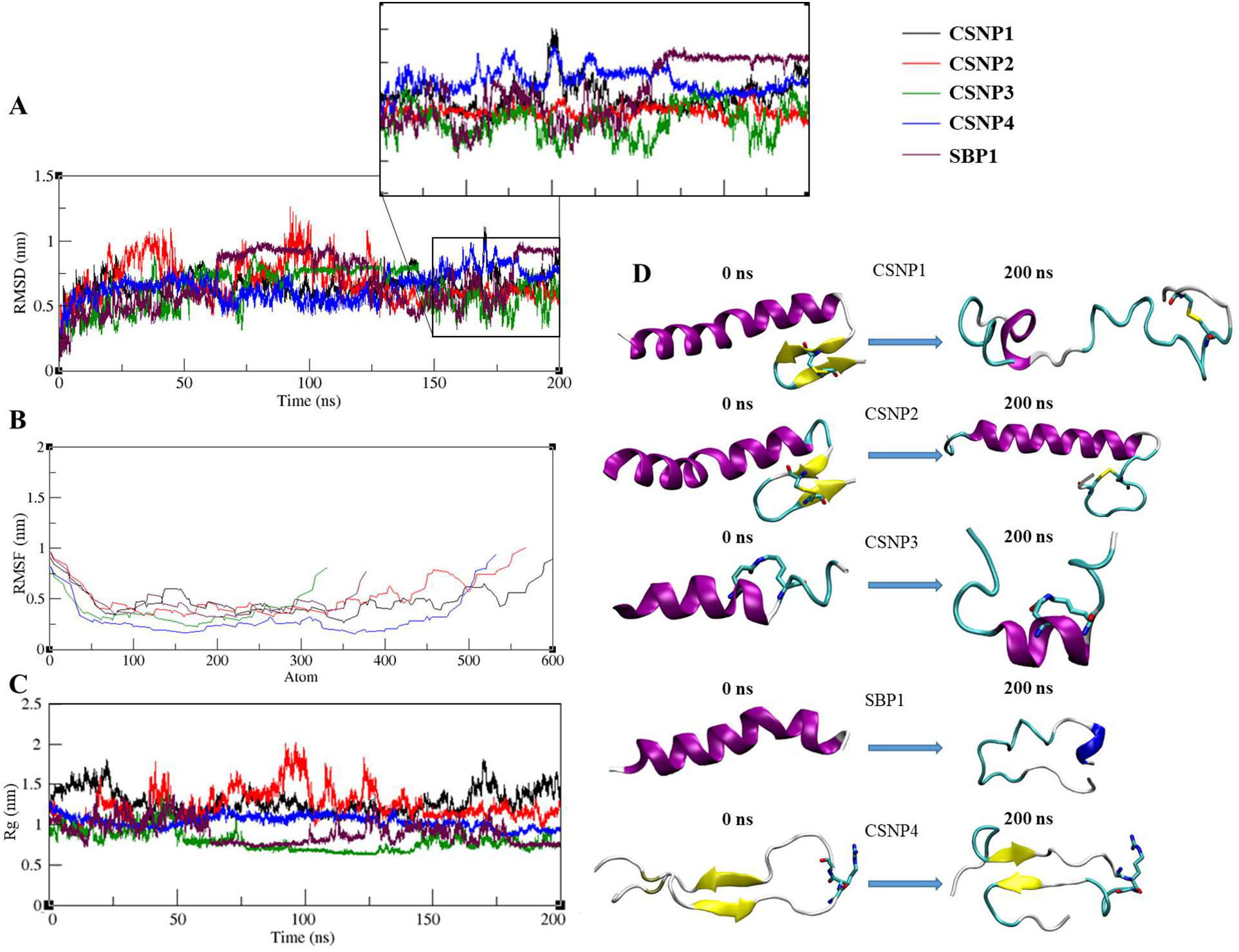
Structural validation of the SARS-CoV-2 spike neutralizing peptides (CSNPs) using molecular dynamics simulations (MDS). **A**) Root mean square deviation (RMSD) plots of all atoms in CSNPs and SBP1 with respect to their initial conformation are presented. The color cods are shown on the top-right and are same of all three (A, B, C) plots. **B**) The root mean square fluctuation (RMSF) plots of the peptides at atomics level. **C**) The radios of gyration of the peptides as a function of time. **D**) The representative 3D structures of the CSNPs and SBP1 peptides showing a significant shift in their structure, as they evolve during simulation. The structural coordinates were taken from the first and last nanosecond of the MDS trajectory. The purple color represents helicity, cyan represent loop and yellow color corresponds to beta sheets. CSNP1 and CSNP3 undergone through a drastic shift in structure. The real-time change in the secondary structures are shown in supplementary movie S1.

### Binding stability of CSNPs to RBD and ACE2

ACE2-derived CSNPs, docked onto RBD, were simulated in a neutralized solution state, whereas CSNP4 was simulated with ACE2. The interface residues of the docked CSNPs were identified and found overlapping with that of ACE2-RBD (**Figure 3A, Supplementary tables 2, 3**). We found that all the target-bound forms of these peptides were relatively more stable compared to their unbound isolated states (**Figure 3B**). However, CSNP3 exhibited an increased RMSD, which might be due to the N-terminal hydrophilic Glutamic acid, causing detachment of the RBD during simulation (**Supplementary movie 2**). SBP1 also affects the overall RMSD of RBD. Indeed, both SBP1-bound RBD and CSNP3-bound RBD had similar tendency in their RMSD plots (**Figure 3C**). To measure the dissociation of CSNPs from their targets, the average distances between the center of their masses and intermolecular hydrogen bonds number were calculated as function of time (**Figure 3D, E**). The average distance between CSNP1-, CSNP2-, and SBP1-RBD complexes remain constant through the simulation course; however, the distances between CSNP3-RBD and CSNP4-ACE2 remained unstable (**Figure 3D**). The distance between ACE2 and CSNP4 was ~35nm at the start of simulation which increased to ~40nm at the midpoint (50ns) of MD run. This increase in the distance could be due to the free N- and C-terminals of the CSNP4, which also had a significant impact on the overall stability (RMSD) of the peptide as well as CSNP4-ACE2 complex (**Figure 3B, C**). The distance between CSNP3 and RBD fluctuated due to the loosely bound hydrophilic N-terminal Glutamic acid of the peptide, which also effect the adjacent Asparagine and detach from RBD (**Supplementary movie 2**). This detachment compels the N-terminal of the peptide on a whip-like motion; nonetheless, the C-terminal residues remained intact with RBD.

**Figure 3:**
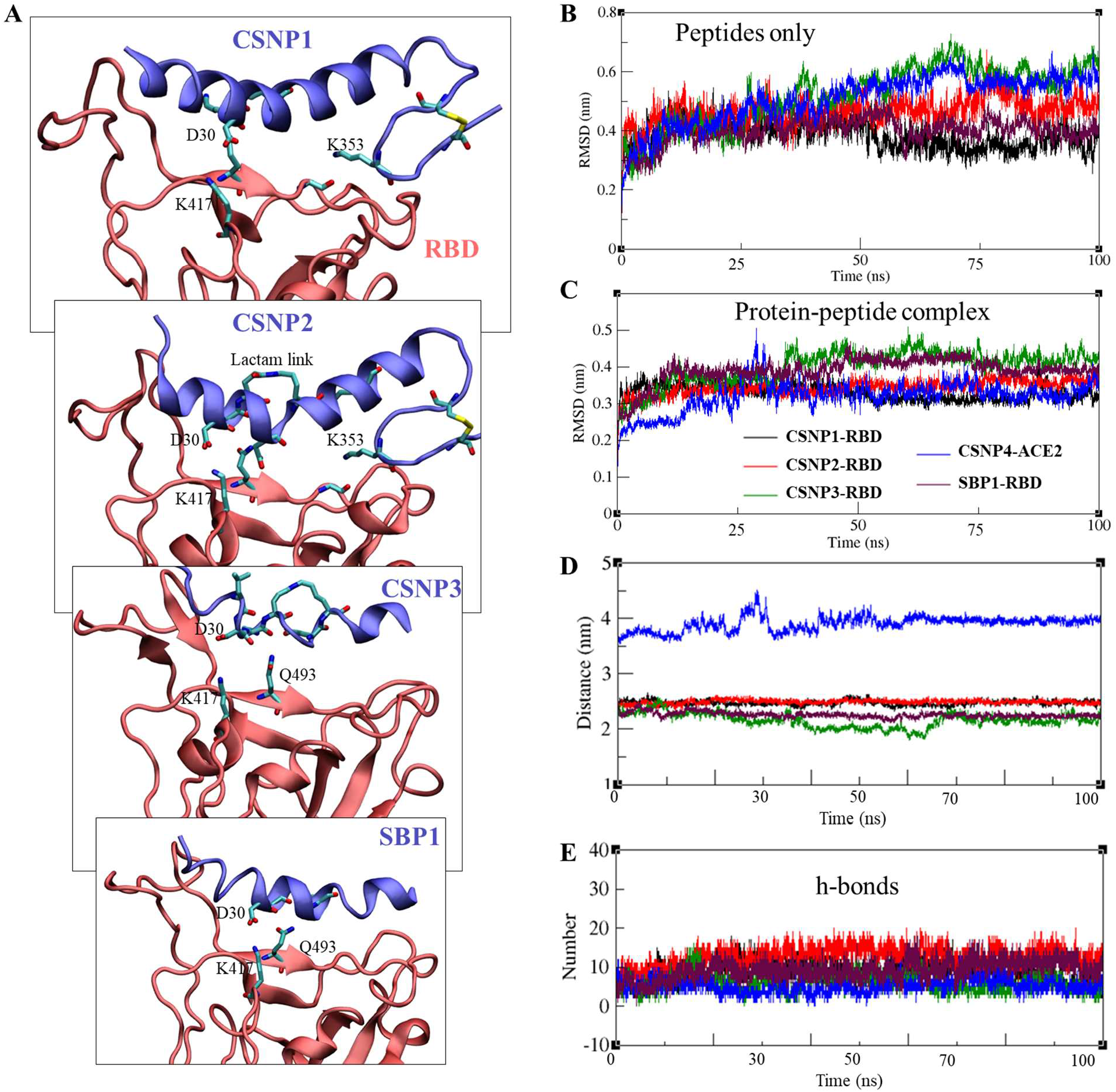
The binding and structural dynamics of CSNPs with SARS-CoV-2 spike RBD and hACE2 as determined through molecular dynamics simulation (MDS). **A**) The initial poses of the helical peptides bound to RBD are shown here. These complexes were used for molecular dynamics simulation. The exact dynamic motions of all complexes during simulation are preserved in supplementary movie S2. **B**) All atoms deviation from their initial position were determined for the peptides through RMSD as a function of time. Peptides acquire a stable dynamics behavior upon binding with their targets, as compared to their isolated solution states (Figure 3A). Overall, there is a gradual rise in the RMSD of CSNP3 and CSNP4. **C**) Among the helical peptides, CSNP1-3 and SBP1, CSNP3 and SBP1 effect the overall stability of the RBD. Although stable, their RMSD plots were distinct from other complexes. CSNP3-RBD in particular exhibit a gradual rise in the RMSD, similar to that of isolated peptide. **D**) The average distances between the center of masses of the peptides and their target proteins. The distances of CSNP3-RBD and CSNP4-ACE2 were not constant as a function of time. **E**) The number of hydrogen bonds established by the peptides with their target proteins as a function of time. The highest number of hydrogen bonds were established and maintained by CSNP2 and RBD. The color cods are same for all plots and given in panel B.

The binding strength between the peptides and their targets were estimated using Poisson–Boltzmann surface area (MM-PBSA) method. For all the five peptide-bound complexes, van der Waals (vdW), Electrostatic (Ele), Polar Solvation (PS), and Solvent accessible surface area (SASA) energies were calculated (**Table 2**). Based on the type and length of the peptides, we could separate them into 3 groups; 1) CSNP1 and CSNP2 contain both the α1 helix and β3-β4 region and binds to RBD; 2) CSNP3 and SBP1 are composed of α1 helix only and bind to RBD; 3) CSNP4 is totally different from the helical peptides and binds to ACE2 and RBD. The overall binding affinities of CSNP1 (total E = −298.44 +/- 82.0 kcal/mol) and CSNP2 (total E = −283.77 +/- 81.3) with RBD were relatively similar. However, the energies of vdW and Ele for CSNP2 were stronger than those for CSNP1 (**Table 2**). Similarly, the total binding energy of CSNP3 helical peptide (total E = −382.73 +/- 63.4 kcal/mol) with RBD was relatively stronger than that of SBP1 (total E = 356.73 +/- 75.1). The polar solvation energy of the SBP1-RBD (853.42 +/- 116.0 kcal/mol) was considerably higher than that of CSNP3-RBD (532.67 +/- 190.2). This notion suggests that upon exposure to solvent, the SBP1-RBD complex may dissociate faster as compared to CSNP3-RBD.

**Table 2:** Binding free energy of the CSNP-bound ACE and RBD complexes.

| Complex | VDW E | Ele E | PS E | SASA E | Total E |
| --- | --- | --- | --- | --- | --- |
| **CSNP1/RBD** | -294.49 +/- 26.0 | -685.26 +/- 74.0 | 719.22 +/- 125.0 | -37.91 +/- 3.3 | -298.44 +/- 82.0 |
| **CSNP2/RBD** | -302.45 +/- 20.3 | -705.56 +/- 96.1 | 763.25 +/- 119.8 | -39.01 +/- 2.4 | -283.77 +/- 81.3 |
| **CSNP3/RBD** | -225.15 +/- 26.6 | -660.4 +/- 159.4 | 532.67 +/- 190.2 | -29.85 +/- 4.3 | -382.73 +/- 63.4 |
| **SBP1/RBD** | -282.12 +/- 23.7 | -889.03 +/- 74.0 | 853.42 +/- 116.0 | -39.00 +/- 2.4 | -356.73 +/- 75.1 |
| **CSNP4/ACE2** | -217.30 +/- 29.6 | -1124.5 +/- 72.6 | 422.46 +/- 135.8 | -27.72 +/- 3.4 | -947.06 +/- 104.2 |

VDW: van der Waal; Ele: Electrostatic; PS: Polar Solvation; SASA: Solvent accessible surface area; E: Energy. All energies are measured in kcal/mol.

### CSNPs interfere with the S1 binding to hACE2

Next, we synthesized the CSNPs to validate their functions as discussed in methods. Unfortunately, we failed to synthesize CSNP1 and excluded in the following biophysical experiments. The human ACE2 (hACE2) overexpressing cells were prepared by transfecting HEK293 cells with hACE2-expressing plasmid, pcDNA3.1-hACE2 and treated with the peptides (**Figure 4A**). We observed that S1 localized to the cell membrane in CSNP-untreated cells, whereas CSNP2 and CSNP4 but not CSNP3 completely abolished the S1-ACE2 interaction (**Figure 4B**). Similar effect was observed when hACE2-HEK cells were treated at higher concentration (25μM) of the peptides (**Supplementary figure 2**). This might be due to the shorter length of CSNP3 which may not be able to optimally bind the RBD and hence fail to abrogate the S1-ACE2 interaction. To validate the S1 localization to the cell membrane and confirm the inhibitory effect of the peptides further, we repeated the experiment in the β-catenin labeled hACE2-HEK cells. As expected, CSNP2 and CSNP4 fully blocked the membrane localization of S1 (**Figure 4C)**.

**Figure 4:**
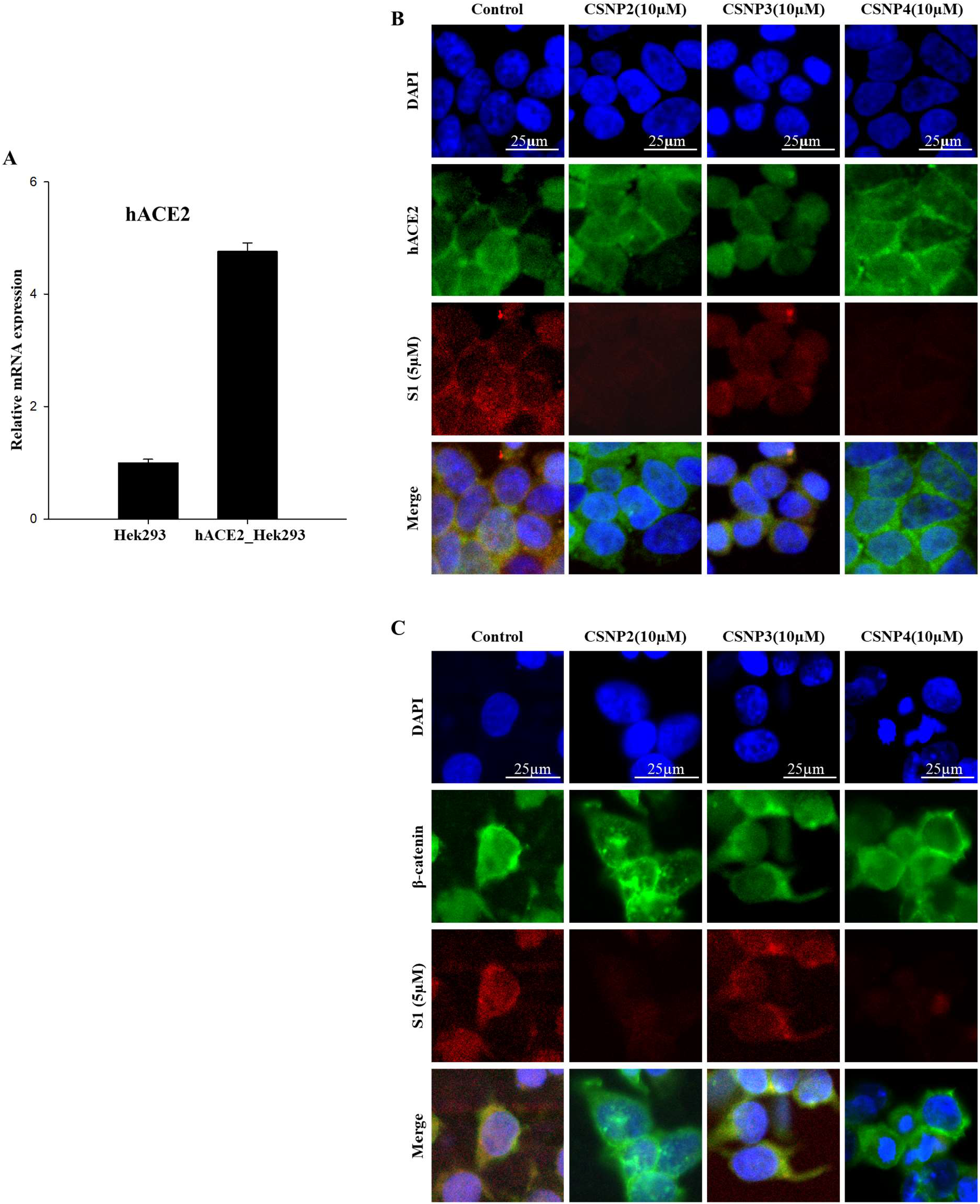
CSNPs peptides deter the SARS-CoV-2 S1 and hACE2 interaction in hACE2-overexpressing HEK293 cells. **A**) The over expression of hACE2 in HEK293 cells as determined through the relative ACE2 mRNA level. **B**) CSNP2 and CSNP4 block the binding of S1 to hACE2 (labeled green) expressing cells. **C**) The membrane localization of hACE2 and the S1 binding to the hACE2 was indirectly confirmed through β-catenin (a protein localizes to adherens junctions) labeling. CSNP2 and CSNP4 clearly deter the membrane binding of S1 protein.

### Biophysical interaction of the CSNPs with target proteins (SPR)

SPR is considered as a reproducible and sensitive technique (33) compared to Biolayer interferometry (34) that can confirm the binding kinetics and biophysical interactions of CSNPs with their targets. Nonetheless, the latter has been also used in the recent biophysical interaction analysis of SARS-CoV-2 spike and multiple ligands including monoclonal antibodies, peptides, and receptor proteins (35). As a validation step, the ACE2 and S1 subunits were immobilized to the CM5 sensor chip as ligands and cross-tested as analytes. Although in nanomolar range, the equilibrium dissociation constant (K_D_) was slightly different in both cases. When S1 was immobilized (as ligand) to the chip the K_D_ was 3.66 nM; whereas, when ACE2 was used as ligand and S1 as analyte the K_D_ was 17.42 nM (**Figure 5A**).

**Figure 5:**
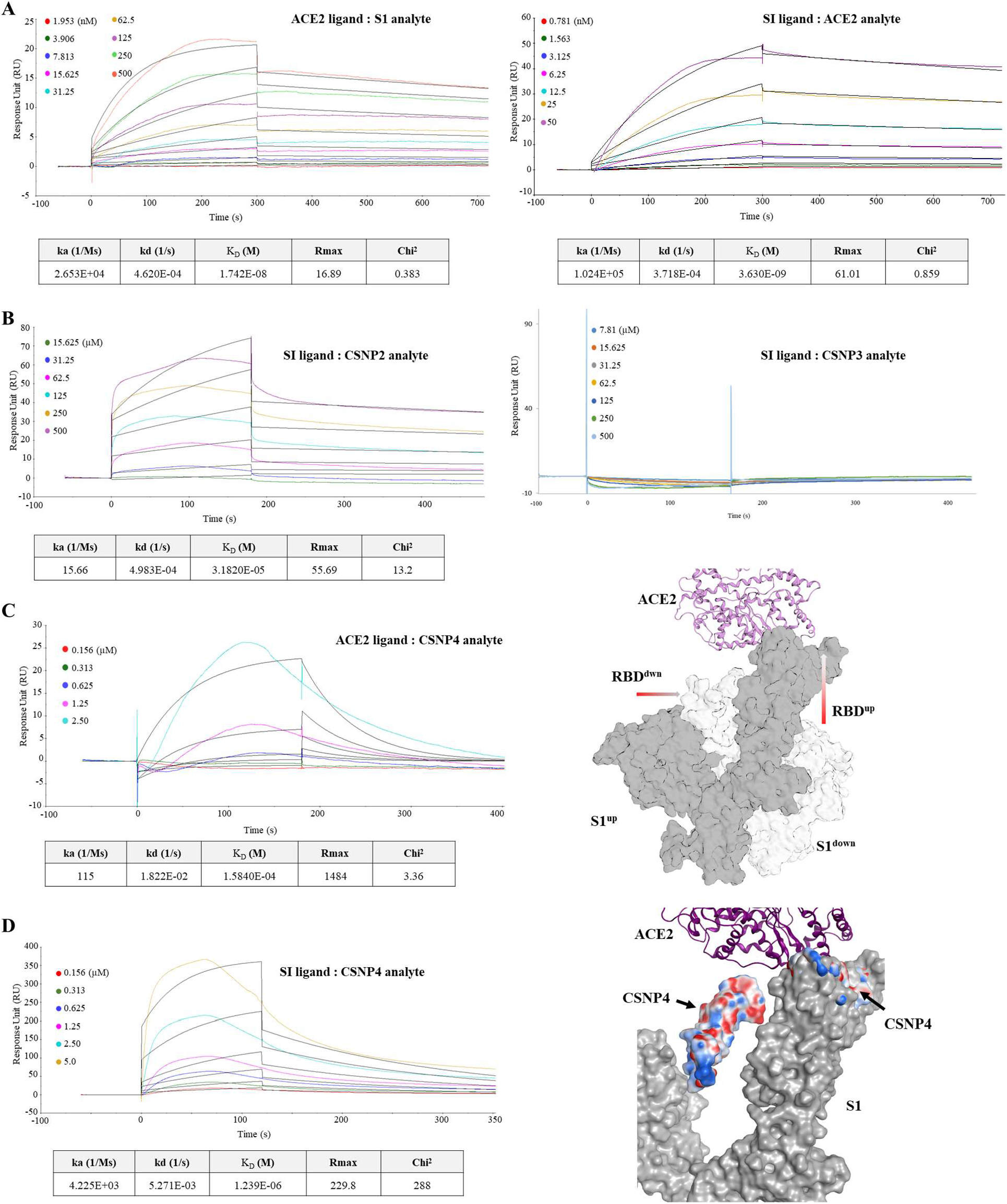
The binding kinetics of the hACE2, SARS-CoV-2 S1, and CSNP peptides. **A**) The binding affinities of hACE2 to SARS-CoV-2 S1 domains and vice versa, were determined by immobilizing both proteins to the chip in separate channels and also used as analytes for injection in different concentrations. The sensorgrams showing 1:1 binding are shown for both proteins and their rates of association (ka), dissociation (kd) and equilibrium constants of dissociation K_D_ are given in the tables below each panel. **B**) Helical peptide CSNP2 binds the SARS-CoV-2 S1 protein dose-dependently, but CSNP3 could not show any binding at the given concentrations. **C, D**) CSNP4 exhibit binding to both hACE2 and SARS-CoV-2 S1 dose-dependently. Nonetheless, the CSNP4-S1 binding affinity was ~13 times higher than that of CSNP4-hACE2. The K_D_ values are given in the tables below each panels and the targets-binding mechanism of CSNP4 are depicted in the cartoon representation.

CSNP2 and CSNP3 were injected as analytes onto the immobilized-S1 at six and seven different concentrations, respectively. CSNP2, but not CSNP3, exhibited a dose-dependent binding to the S1 protein with K_D_=31.8 μM (**Figure 5B**). This finding suggests that CSNP2, and probably CSNP1, retains its binding to RBD. S1-ACE2 binding was relatively stronger (K_D_= 3-17 nM) compared to the CSNP2-RBD binding. This may confirm that in addition to the α1 helix and Lys353, auxiliary residues of the ACE2-RBD interface further enhance the binding (**Figure 1A, Table 1**). Further studies might be required, including the X-ray diffraction analysis of CSNP2-RBD that could pave a way for the structural guided molecular modification CSNP2 to enhance its binding and specificity to RBD.

Next, we measured the binding kinetics of CSNP4 and immobilized ACE2, revealing relatively weak and dose-dependent binding-affinity between them (K_D_=158 μM) (**Figure 5C**). Considering our immunocytochemistry results (**Figure 4)** and the fact that CSNP4 occupies the junction between the NTD and RBD^up^ of the adjacent S protomer in the trimeric Spike protein (**Figure 5C**), we evaluated the binding affinity of CSNP4 with S1. Unexpectedly, we found that CSNP4 binds S1 as strong as CSNP2. The weak binding affinity of CSNP4-ACE2 can be explained for two reasons. First, the α1 helix of ACE2 provides a narrow and shallow surface for the CSNP4 binding. Second, Lys417, which contributes significantly to the ACE2-RBD binding (**Table 1**), is not included in CSNP4. From this data, we suggest that designing an RBD-derived decoy peptide against ACE2 may not be sufficient to block ACE2-RBD interactions due to the large conformational space and irregular loop structures of the ACE2-bindng motifs in RBD. The CSNP4-based S1 neutralization could be attributed to the stronger CSNP4-S1 binding, which may limit the freedom of RBD and ultimately deter it’s binding to ACE2.

## Discussion

Hots cell entry is the first step in viral infection and remained a priority concern for the therapeutic interventions. The impulsive conformation switching of the SARS-CoV-2 RBD, as reported by Wrapp et al. (15), is one of the immune evasion strategy utilized by CoVs (11, 13). After understanding the receptor binding mechanism of SARS-CoV-2, we have tried to explain its immune evasion mechanism through a stepwise illustration (**Figure 6A**). ACE2 is the principal RBD-recognizing receptor, which exists in both membrane-bound and soluble form. ADAM17 (a disintegrin and metalloproteinase 17) (36) and TMPRSS2 (37) regulates the shedding of its enzymatically active ectodomain into the blood circulation (**Figure 6B**). Watson et al. have demonstrated that soluble ACE2 binds the RBD with comparatively similar binding affinity as do its neutralizing antibodies (38). In fact, the soluble ACE2 serves as “double edge sword”; on one hand it acts as a potent inhibitor of the pseudotyped lentivirus *in vitro*(39), on the other hand it abrogates the RBD neutralization (**Figure 6A**). Due to its RBD-binding ability, soluble ACE2 has been used to block viruses from entering cells (40). Collectively, RBD spontaneously switches from ‘up’ to ‘down’ and that soluble ACE2 shields the spike protein from immune surveillance. This paradox should be considered for the identification of therapeutic agents that abolish the Spike-ACE2 interaction. Instead of alleviating certain symptoms, a world-wide affordable and deployable strategy need to be implemented to design stable and frequently administrable therapy that could offset the real cause of infection. Biologics that can interfere with RBD-ACE2 binding, facilitating the “up” conformation of RBD, and minimally interfere the antibody binding and B cell response to the S protein are required. This strategy was partly utilized by the two FDA approved RNA-based vaccines, BNT162b2 and mRNA-1273, that encode the SARS-CoV-2 spike protein in a stable prefusion conformation. In both vaccines the antigenic spike protein is mutated by substituting the two consecutive lysine 986 and valine 987 at the top of central helix in S2 domain into proline to convert the metastable prefusion conformation into a stable prefusion state for continuous immune surveillance (1, 41, 42).

**Figure 6:**
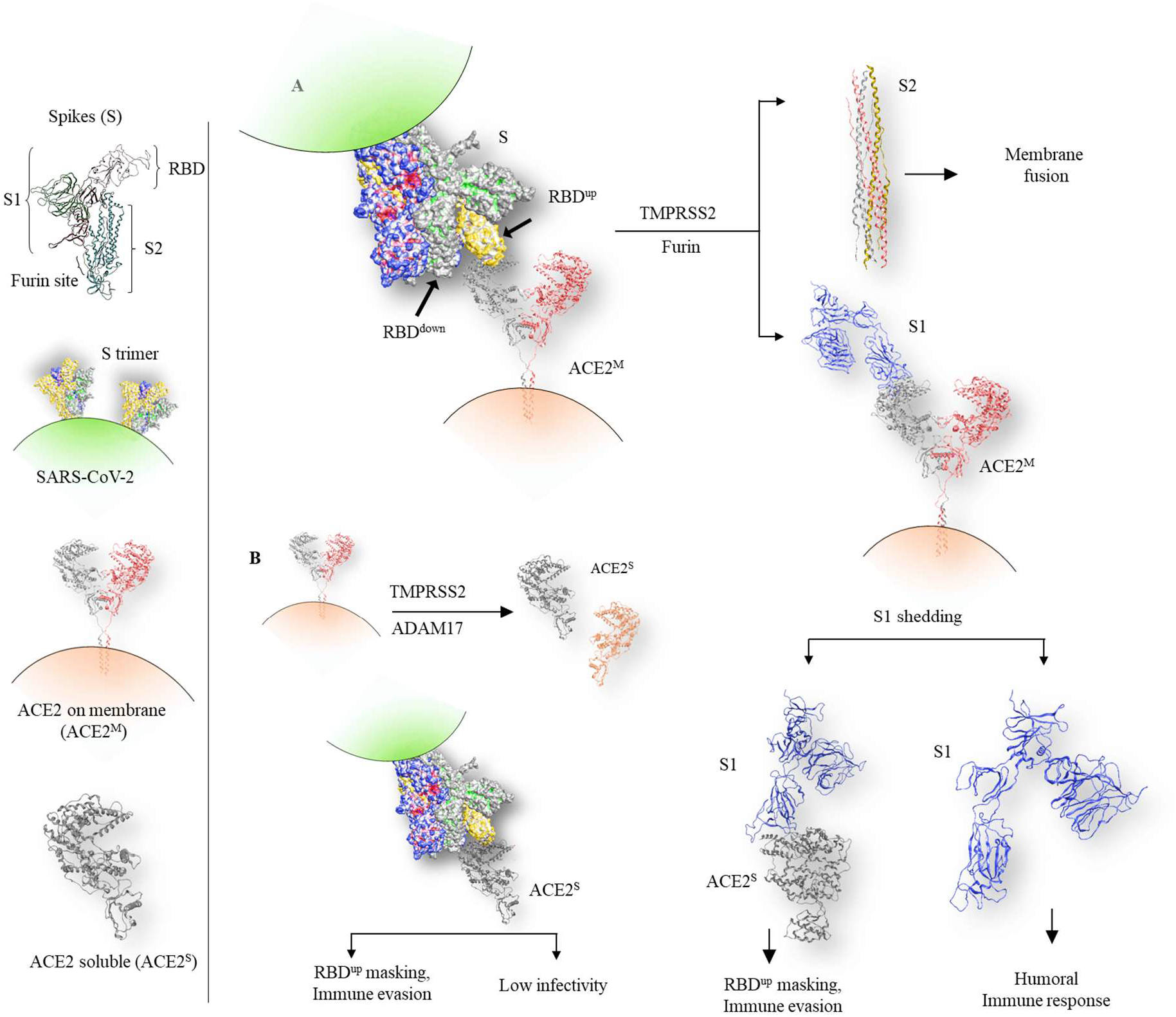
The ACE2-binding and shedding mechanism of SARS-CoV-2 spike protein S1 subunit. **A**) The receptor binding domain (RBD) in S1 subunit of the spike protein binds the membranebound ACE2 (ACE2^M^). The cell surface expressed and lysosomal proteases, including cathepsin, furin, and TMPRSS2 cleaves the ACE2^M^-bound spike at the S1-S2 cleavage motif. S2 subunits then facilitate the viral and host membrane fusion while S1 subunits are shed extracellularly. These stranded S1 subunits are immunogenic and crucial for host antiviral humoral immune response. The released S1 subunits could be masked by the overexpressed soluble ACE2^S^ protein in diabetic and cardiac patients, causing poor viral neutralization and comparatively high mortality. **B**) The soluble ACE2 (ACE2^S^) or its mimics could neutralize and deter the binding of spike to membrane-bound ACE2. This could cause a paradoxical condition of low viral infectivity yet low viral neutralization by host antibody response.

The binding spike with ACE2 and the subsequent conformational rearrangements of the S2 for the viral-host membranes fusion make this entire process a prime target for the vaccines and drug development. We and others have investigated that SARS-CoV-2 utilize shallow and expanded surface contacts between spike and ACE2 for host cells entry (15, 18, 31, 35). These hurdles i.e. the shallow, widely expanded, and flat contact surfaces, makes the spike-ACE2 a hard target for widely utilized small organic molecules-based medicinal chemistry approaches. Even though, tremendous progress has been made in targeting viral protease with small molecules (43), disrupting spike-ACE2 interface with same strategies remained an uphill task. Alternatively, small peptides, peptide-memetic, and mini-proteins have overcome this difficulty and we have seen astonishing outcomes in blocking viral entry and neutralization (26, 44, 45). The deployment of peptides-based biologics has expanded the concept of druggability by specifically and effectively targeting PPI that are hard to target with organic molecules (46). Despite the considerably high selectivity and lower toxicity, compared to small molecules, peptides therapeutics are challenging due to their compromisable stability and subsequent low bioavailability due to shortened half-lives in vitro and in vivo. Improved strategies including peptides stapling, lipid and polyethylene glycol (PEG) modification, and peptide bundles have been advised over the time to overcome these hurdles and formulated effective biologics against SARS-CoV-2 spike protein (26, 35, 47).

Similarly, CSNPs were designed to block the ACE2-RBD interaction, interfere with capping of spike by soluble ACE2, and present spike proteins in its open form to host B-cell responses. The helical peptides, CSNP1-3, were designed to be engaged in the binding of spike to ACE2. A number of studies have reported redundant yet overlapping epitopes on the SARS-CoV-2 RBD that are recognized by neutralizing antibodies (48–50). Among them, Ju et al. isolated 206 RBD-specific monoclonal antibodies from 8 SARS-CoV-2 patients, which compete with ACE2 for RBD binding (49). After structure superimposition of the P2B-2F6-RBD (isolated by Ju et al.) and CSNP1-RBD, we could observe that CSNP1 was not able to make any stearic hindrance with P2B-2F6 (**Supplementary Figure 1B**). This suggests that CSNP2, which binds the same RBD-interface as CSNP1, hinder the ACE2-RBD interaction (**Figures 4 & 5**) and allow RBD to be recognized by neutralizing antibodies. However, this notion needs further confirmation through competitive binding assays. Second, the immune-evasive position-switching of RBD as well as its ACE2 binding require a strategy to restrict the freedom of RBD for robust immune response, at the same time hinder its binding to ACE2. As a matter of fact, RBD itself could be used as double-edge sword in this scenario. In its closed conformation (RBD^down^), the ACE2-binding motifs of RBD occupy the “V” shaped space between the NTD and RBD of the adjacent S protomer (**Figure 5C, D**). Consequently, CSNP4 was designed and found to block the S1 binding to hACE2 (**Figure 4B**). We believe that CSNP4 not only inhibits ACE2-RBD binding but can also compete with RBD hiding in the “V” pocket.

The three dimensional conformational spacing of the pharmacophores in α-helix is crucial for the helical CSNP1-3 peptides. Short peptides unfold and lose their secondary structures in solution state when they were truncated from tertiary folded proteins. The structural stability of the CSNPs investigated through MDS indicates that the unconstrained peptides very quickly loses their helicity and acquire irregular looped conformation (**Figure 2D, Supplementary movie S1**). This may be one of the reasons short peptides lose target specificity. To overcome this limitation, two research groups have stabilized the α1 helix by increasing the helical bundles and designed peptide biologics that could effectively inhibit SARS-CoV-2 cell entry (44, 45). However, this modification increases the size of these peptides by ~3-4 folds, increasing the cost of synthesis and formulation. We and others have previously used peptide stapling to enhance the target specificity of therapeutic peptides (28, 51). In fact, Fiarlie and his co-workers found that the helical-constrained compounds hold comparatively similar biological potencies in PPI as their parent proteins. They constructed four such compounds and proposed that downsized-constrained peptides could be of great value in biological PPIs and medicine (52). Unfortunately, we could not synthesize the CSNP1 to compare its potency with CSNP2; however, CSNP3 (a relatively shorter version of CSNP1) was unable to bind SARS-CoV-2 S1 (**Figure 5**). This notion suggests that in addition to the structural integrity, optimum length and the featured pharmacophores are vital for CSNP to bind RBD. Together, we suggest that structure guided computational medicinal chemistry and click chemistry approaches might be useful in designing CSNPs to enhance tethering to their targets. Collectively, we suggest that CSNP2 and CSNP4 are stable peptide that deter the binding of ACE2 and S1 subunit of SARS-CoV-2 and could be potent antidote for COVID-19.

## Methods

### Peptides Designing Rationale and Synthesis

The crystal structure of SARS-CoV-2 spike-RBD bound to ACE2 (PDB ID: 6M0J) was used to design the CSNP1-CSNP3 peptides. For CSNP4, both the trimeric spike-ACE2 (PDB ID: 6ZXN) and ACE2-RBD complexes were considered. For CSNP1-3 the hotspot residues on both RBD and ACE2 were designated through PDBePISA (53) and their contribution into the interface were evaluated through alanine scanning, using DrugscorePPI (54). This server utilizes the interface knowledge and calculate the difference in binding energy of the wild type (ΔG^WT^) and mutant (ΔG^MUT^) residues at the interface and provides hotspot information in terms of numerical values and the corresponding 3D b-factor coordinates. Two regions on the ACE2, a.a. 23-46 and a.a. 352-357 that optimally engage the RBD, were selected for the parent peptides (CSNP1) design. Both regions were linked through GPG loop and the freedom of K353 was restricted by disulfide bond (S-S) bond, stapling the two beta sheets at position C350 and C356.

The electrostatic surface map, around the hotspot and other interface residues were created using APBS and APBSrun plugins and potential modifiable and vital pharmacophores were identified in the α1 helix (a.a. 23-46) using pharmacophore package in MOE (2019.0102). Taking the surface complementarity into consideration, five potential residues, i.e. Glu23, Lys26, Thr27, His34, and Gln42 were identified and subsequently substituted to enhance the CSNP1-RBD binding using residues-scan tool in protein-design package of MOE suit. The detailed protocol has been outlined previously (55). In first step, the mutation window was restricted to one residue only, and the Glycine and self-mutation were excluded during mutant generation. Based on binding affinity and stability, the top five mutants were selected and subjected to the second round of residues scan, keeping the mutation window five. Based on binding Change in the binding affinity of the wild type and mutant peptides were dually evaluated through SSIPe and EvoEF (56). To stabilize and retain the helical structure of the selected peptides a lactam bridge was created at i and i+4 position of the non-interface residues, as described in our previous work (28). For CSNP3, the pharmacophores in the α1 (a.a. 21-46) region of ACE2 were considered but the a.a. 352-357 regions were excluded. The CSNPs designing strategy has been outlined in figure 2C. Like CSNP1, CSNP4 was designed by linking a.a. 445-456 and a.a. 488-501 of the SARS-CoV-2 RBD and residues scan was not implemented. Three, CSNP2-4, of the final selected CSNPs peptide were successfully synthesized by Peptron Inc. (Daejeon, Korea) at a purity of 99% (CSNP2), 95% (CSNP3), and 96% (CSNP4) as determined by reversed-phase high-performance liquid chromatography (HPLC; Shimadzu Prominence). Detailed protocol has been provided in our previous study (28).

### Molecular Dynamics Simulation (MDS) analysis of CSNPs

MDS is widely used technique to study the protein folding and the dynamic behavior of macromolecules in complex or isolated form. Briefly, default ABMER99-ILDN force-field (57) was used for CSNP1, CSNP4, and SBP1 (a previously identified RBD-binding peptide) simulation, while the same force-field was modified for the lactam-stapled peptides, CSNP2 and CSNP3. New residues and parameters were added into the modified force-field wherever needed. The peptides in their isolated form were simulated for 200 ns and their target-bound complexes for 100 ns. All simulations were carried out in GROMACS 2019.6 in TIP3P water filled cubic box with 10 Å extended boundaries from the protein. All the systems were neutralized with counter ions, Na^+^/Cl^-^, wherever needed and energy minimized under steepest descent algorithm, then equilibrated with NVT ensemble for 0.2 ns and re-equilibrated with NPT ensemble for 0.2 ns, under constant temperature and pressure, respectively. The temperature and pressure were coupled with V-rescale and Parrinello-Rahman barostat methods (58), respectively. The bond lengths were constrained with LINCS algorithm and long-range electrostatic interactions were computed with particle mesh Ewald algorithm (59).

### Binding free energy calculations using Molecular mechanics Poisson-Boltzmann surface area

Molecular mechanics Poisson-Boltzmann surface area (MM-PBSA) is used for calculating the relative biding energy of ligands bound to target (60). We used the g_mmpbsa and APBSA tools implemented in GROMACS for calculating energies. As g_mmpbsa tool is compatible with older versions of GROMACS (versions 5 or lower), the “tpr” files created by GROMACS 2019.6 were recreated through GROMACS 5.1 and used for binding energy calculation. The relative binding energies of the complexes were approximated according to the following energies terms.

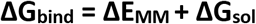

whereas;

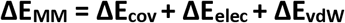

and;

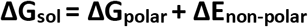

ΔE_MM_ is gas-phase MM-energy change and ΔG_sol_ is the solvation free energy change. van der Waals energy change (ΔE_vdW_), the electrostatic energy change (ΔE_ele_), and the covalent energy change (ΔE_cov_). The solvation free energy (ΔG_sol_) is computed by combining both polar and non-polar energies. All these changes were computed via ensemble which is averaged over a set of conformations sampled over the last 25 ns simulation trajectory at 0.01 ns time interval.

### Computational tools used in this study

For simple visualization and collecting structural insights of the SARS-CoV-2 spike and ACE2 proteins, free available packages of VMD (61), Pymol (https://pymol.org), and Chimera (62) were utilized. For electrostatic surfaces isolation of the proteins, APBS and APBSrun plugins in Pymol and VMD, and for monitoring changes in the secondary structures of the peptides as function of time the sscache.tcl script was used in VMD. For 3D animated movies creation VMD was used. For the interface analysis and determining the contribution of each residue into the ACE2-S binding, online server PDBePISA (v1.52)(53) and free available BIOVIA Discovery Studio Visualizer were used. After using the PPCheck hotspot prediction tool, the alanine scanning package in same sever was used for the aniline mutagenesis (63). The hotspot results were validated through DrugScorePPI web server and the results were recorded in terms of energies (54, 64). For pharmacophore evaluation ligandScout trial version and MOE were used (65). However, as our interest lays in amino acid determinants isolation, we did not use the pharmacophore model for drug screening. For molecular dynamics simulations GROMACS 2019.6 was used (66). For MM-PBSA calculations, the “tpr” files created by GROMACS 2019.6 were recreated through GROMACS 5.1 and used for binding energies calculations, as described previously (64).

### Surface Plasmon Resonance (SPR) analysis

For the physical interaction of CSNPs with ACE2 and S1 subunit of SARS-CoV-2, SPR assay was conducted, using Biacore T200 (GE Healthcare, Sweden) technique. S1 (ligand, AcroBiosystems, S1N-C52H3-100UG, USA) protein was immobilized to the CM5 sensor chip (GE Healthcare, Cat#. BR-1005-30) at 6.0 μg/mL concentration using 10mM sodium acetate (pH 5.5) as immobilization buffer. ACE2 (Acrobiosystems, AC2-C52H7-50ug, USA) was immobilized to the same chip at 6.2 μg/mL concentration using 10mM sodium acetate immobilization buffer. The following solutions were used as running buffers: 1) HBS (10mM HEPES, pH 7.4, 150mM NaCl), 2) HBS-EP (10mM HEPES, pH 7.4, 150mM NaCl, 3mM EDTA, 0.05% P20), 3) HBS with 5% DMSO. NaOH solution (10 ~ 50) mM was used as regeneration buffer. The CSNPs were injected into the ligand-bound chip at different concentrations and their respective k_a_, k_d_, and K_D_ values were calculated. The running buffer was injected into the empty channel as a reference. The experiments were conducted in duplicate with freshly prepared reagents, and the data were analyzed in the Software Control (version 2.0.1), and BIAevaluation (version 3.0) software.

### Immunocytochemistry and confocal microscopy analyses

HEK293 (human embryonic kidney 293) cells were purchased from the Korean Cell Line Bank (KCLB, Chongno, Seoul, Korea) and cultured on growth media consisting of Dulbecco’s modified Eagle’s medium (DMEM, Gibco, USA) with 10% fetal bovine serum (FBS, Gibco, USA) and 1% antibiotics (Gibco, USA) in a 5% CO2-humidified incubator. For Immunocytochemistry analysis, HEK293 cells were transfected with 3.0ug pcDNA3.1-hACE2 (Addgene, 145033, USA) plasmids by Lipofectamine 3000 (Invitrogen, L3000015, USA). The relative hACE2 quantity was evaluated through mRNA expression level using following primers: (Cosmo genetech, hACE2-F: 5’-TCC ATT GGT CTT CTG TCA CCC G-3’, hACE2-R: 5’-AGA CCA TCC ACC TCC ACT TCT C-3’, Republic of Korea). For CSNPs-binding, hACE2-overexpressed Hek293 cells were incubated with 10 uM peptide for 1 hour and then treated and incubated with 5 uM SARS-CoV-1 S1 protein-His Tag (AcroBiosystems, S1N-C52H3-100UG, USA) for 24 hours. After the cells were washed three times with PBS, incubated with serum free media with primary antibody anti-ACE2 (1:100, Cell signaling, 4355S, USA), anti-CTNNB1 (1:100, Cell signaling, 8480S, USA) and anti-His-Tag (1:100, Santa cruz, sc-8036, USA) at 4°C for 2 hours. Cells were then fixed with 4% paraformaldehyde for 5 min and incubated with donkey anti-mouse IgG conjugated with Alexa Fluor 594 (1:200, Thermo, A21203, USA) and donkey anti-rabbit IgG Alexa Fluor 488 (1:200, Thermo, A21206, USA) at room temperature for 2 hours. Before the secondary antibodies incubated, cells blocking with blocking solution (1% PBS containing 1% BSA and 0.1% Tween 20) at room temperature for 1 hour. All secondary antibodies were diluted in an appropriate concentration of blocking solution. Nuclei were stained with DAPI containing mounting solution (Vector, H-1200, USA). The cells were then visualized on a LSM710 (Carl Zeiss) confocal microscope.

## Acknowledgements

This research was supported by grants from the National Research Foundation of Korea (NRF) funded by the Ministry of Science and ICT (MSIT) (NRF-2017M3C9A6047620, NRF-2019R1A5A2026045, NRF-2017M3A9B6061509), Republic of Korea. This study was also supported by KREONET (Korea Research Environment Open NETwork) which is managed and operated by KISTI (Korea Institute of Science and Technology Information).

## Authors contributions

M.S. and H.W. contributed toward conceptualization of the project and designed he methodology; M.S. and S. U. M. performed the investigation and formal analysis; M.S., S. U. M., and H.W. wrote the original manuscript draft; H.W supervised the study and provided funding acquisition; all the authors contributed to editing and reviewing the manuscript.

## Competing interests

All authors declare that there is no competing interest

## Notes

### Competing Interest Statement

The authors have declared no competing interest.

