## Supplementary material for "Pharmacophore-based peptide biologics neutralize SARS-CoV-2 S1 and deter S1-ACE2 interaction *in vitro*": Supplementary DATA.docx

**^‡^ Contributed equally**

***Corresponding author**

Fax number: 82-31-219-5049,

**Supplementary DATA**

**Supplementary Tables**

**Supplementary Table 1. Mutations at the five positions in CSNP1 and their resulting binding energy differences with respect to**

|  | **Mutation** | **Absolute Affinity (kcal/mol)** | **Relative binding affinity (kcal/mol)** |
| --- | --- | --- | --- |
| CSNP2 | E23L, K26A, T27N, H34D, Q42K | -612.91 | -2.39 |
| CSNP2-1 | E23L, K26K, T27N, H34H, Q42K | -612.84 | -2.32 |
| CSNP2-2 | E23E, K26K, T27N, H34H, Q42K | -612.37 | -1.84 |
| CSNP2-3 | E23L, K26A, T27T, H34H, Q42K | -612.16 | -1.63 |
| CSNP2-4 | E23E, K26A, T27N, H34D, Q42K | -612.08 | -1.55 |
| CSNP2-5 | E23L, K26K, T27T, H34H, Q42K | -612.07 | -1.54 |
| CSNP2-6 | E23E, K26K, T27T, H34H, Q42K | -612.01 | -1.48 |
| CSNP2-7 | E23E, K26A, T27T, H34H, Q42K | -611.94 | -1.41 |
| CSNP2-8 | E23E, K26A, T27N, H34H, Q42K | -611.89 | -1.36 |
| CSPN2-8 | E23E, K26K, T27N, H34D, Q42K | -611.55 | -1.02 |
| CSNP2-9 | E23L, K26A, T27N, H34H, Q42K | -611.44 | -0.91 |
| CSNP2-10 | E23L, K26A, T27N, H34H, Q42Q | -611.06 | -0.53 |
| CSNP2-11 | E23E, K26K, T27N, H34H, Q42Q | -610.99 | -0.46 |
| CSNP2-12 | E23E, K26A, T27N, H34D, Q42Q | -610.87 | -0.34 |
| CSNP2-13 | E23E, K26K, T27T, H34D, Q42Q | -610.73 | -0.20 |
| CSNP2-14 | E23E, K26K, T27T, H34D, Q42K | -610.66 | -0.13 |
| CSNP1 | E23E, K26K, T27T, H34H, Q42Q | -610.52 | 0 |

**Supplementary Table 2: The Hydrogen bonds and salt bridges at the interface of CSNP-RBD and their bonds-strength in term of energies.**

| **CSNP** | **Peptide** | **RBD** | **Type** | **Dist Å** | **E^kcal/mol^** |
| --- | --- | --- | --- | --- | --- |
| **CSNP1** | Asp30 | Lys417 | IH | 3.12 | -25.1 |
|  | Lys31 | Gln493 | H | 2.71 | -7.6 |
|  | Glu35 | Gln493 | H | 2.75 | -8.9 |
|  | Glu37 | Tyr505 | H | 2.58 | -3.3 |
|  | Asp38 | Gln498 | H | 2.86 | -5 |
|  | Tyr41 | Gln498 | H | 2.95 | -1.2 |
|  | Lys353 | Gly496 | H | 2.76 | -11.5 |
|  | Lys353 | Gly502 | H | 3 | -3.6 |
| **CSNP2** | Gln24 | Asn487 | H | 2.92 | -1.9 |
|  | Asn27 | Tyr473 | H | 2.8 | -2.8 |
|  | Asp30 | Lys417 | IH | 2.71 | -31.6 |
|  | Lys31 | Gln493 | H | 2.69 | -1.3 |
|  | Asp34 | Tyr453 | H | 2.57 | -3.8 |
|  | Glu35 | Gln493 | H | 2.76 | -6.5 |
|  | Glu37 | Tyr505 | H | 2.67 | -4.2 |
|  | Asp38 | Tyr449 | H | 2.62 | -3.8 |
|  | Tyr41 | Gln498 | H | 2.71 | -2 |
|  | Lys42 | Gly446 | H | 2.88 | -10.5 |
|  | Lys353 | Tyr495 | H | 2.93 | -9.8 |
|  | Lys353 | Gly502 | H | 2.99 | -4.2 |
|  | Arg357 | Thr500 | H | 2.97 | -4.6 |
| **CSNP3** | Asn24 | Asn487 | H | 2.82 | -2.3 |
|  | Asn27 | Tyr473 | H | 2.84 | -2.5 |
|  | Asp30 | Lys417 | IH | 2.68 | -35.1 |
|  | Lys31 | Gln493 | H | 2.69 | -8 |
|  | Glu35 | Gln493 | H | 2.72 | -6.6 |
|  | Glu37 | Arg403 | IH | 3.2 | -13.4 |
|  | Glu37 | Tyr505 | H | 2.58 | -5 |
|  | Asp38 | Tyr449 | H | 2.67 | -4.1 |
|  | Tyr41 | Gln498 | H | 2.66 | -3 |
| **SBP1** | Asp30 | Lys417 | IH | 2.74 | -30 |
|  | Lys31 | Gln493 | H | 2.74 | -11 |
|  | Glu35 | Gln493 | H | 2.76 | -5.2 |
|  | Glu37 | Arg403 | IH | 3.06 | -33.7 |
|  | Glu37 | Tyr505 | H | 2.65 | -2.7 |
|  | Asp38 | Gln498 | H | 2.97 | -6 |
|  | Asp38 | Asn501 | H | 2.85 | -8.3 |

**Supplementary Table 3: The Hydrogen bonds and salt bridges at the interface of CSNP4-ACE2 their bonds-strength in term of energies.**

| **CSNP4** | **ACE2** | **Type** | **Dist Å** | **E kcal/mol** |
| --- | --- | --- | --- | --- |
| Asn330 | Trp445 | H | 3.41 | -0.6 |
| Asp38 | Gln447 | H | 2.77 | -5.8 |
| His34 | Tyr453 | H | 2.75 | -1.5 |
| Gln24 | Arg460 | H | 2.84 | -9.5 |
| Thr27 | Arg460 | H | 2.95 | -2.6 |
| Lys31 | Gln493 | H | 2.75 | -11.5 |
| Glu35 | Gln493 | H | 2.8 | -4.7 |
| Lys353 | Tyr495 | H | 2.76 | -4.8 |
| Asp38 | Gly496 | H | 3.56 | -0.7 |
| Lys353 | Gly496 | H | 3.36 | -0.6 |
| Asp355 | Thr500 | H | 2.62 | -1.4 |

**Supplementary Figures**

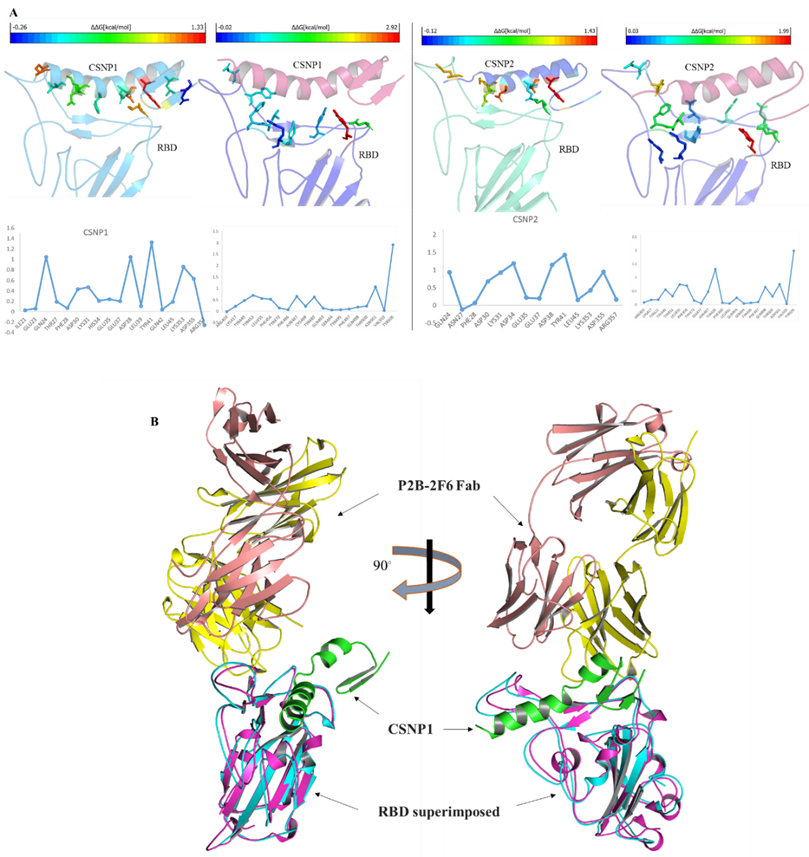

**Figure S1: Hotspots identification at the SARS-CoV-2 spike RBD and hACE2 interface. A**) The hotspot residues on CSNP1 peptide and hACE2 (left panel) are shown in cartoon (colored according to the b-factor) and their binding energy contributing are shown in the plots below. The right side panel show the hotspots on CSNP2 with respect to hACE2. Residues substitution in CSNP2 evenly distribute the hotspots and exhibits stronger affinity as compared to CSNP1. **B**) The SARS-CoV-2 spike RBD-CSNP1 complex and RBD-P2B-2F6 (Fab region of RBD neutralizing monoclonal antibody) are superimposed. The CSNP1 and Fab binding interface in RBD do not overlap.

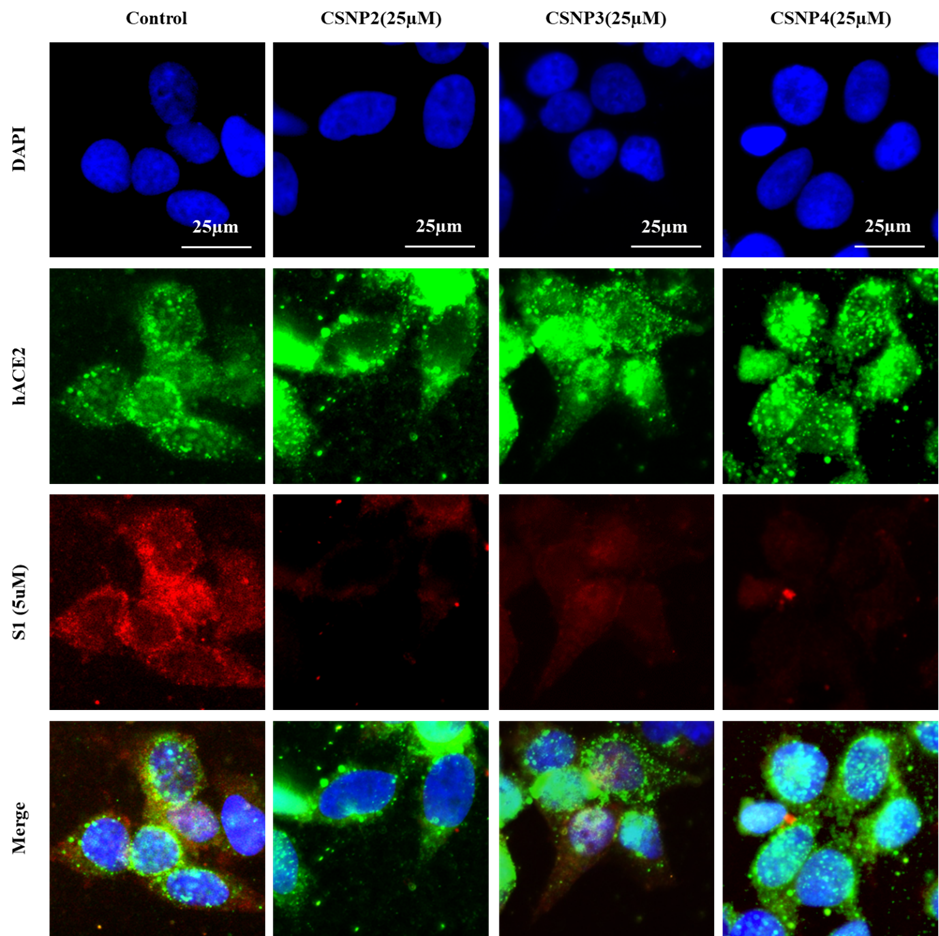

**Figure S2:** CSNPs peptides deter the SARS-CoV-2 S1 and hACE2 interaction in hACE2-overexpressing HEK293 cells at 25µM

**Captions for supplementary Movies**

**Supplementary movie S1:** The 3D motion in this video represents the structural dynamics of CSNP and SBP1 (control) peptides in solvent. The changes in the colors are assigned to track the folding of the peptides. Purple color represents helix, yellow color represents beta sheets, and blue or cyan colors corresponds to a transient turn or loop structure. These 3D movies are based on 1000 structural frames evenly extracted from the molecular dynamics trajectory.

**Supplementary movie S2:** The 3D motion in this video represents the structural dynamics of CSNP and SBP1 (control) peptides bound to their targets. The licorices are shown to track the changes in the interface residues. These 3D movies are based on 1000 structural frames evenly extracted from the 100 ns molecular dynamics trajectory.
